# Collective action and the collaborative brain

**DOI:** 10.1101/009746

**Authors:** Sergey Gavrilets

## Abstract

Humans are unique both in their cognitive abilities and in the extent of cooperation in large groups of unrelated individuals. How our species evolved high intelligence in spite of various costs of having a large brain is perplexing. Equally puzzling is how our ancestors managed to overcome the collective action problem and evolve strong innate preferences for cooperative behavior. Here I theoretically study the evolution of social-cognitive competencies as driven by selection emerging from the need to produce public goods in games against nature or in direct competition with other groups. I use collaborative ability in collective actions as a proxy for social-cognitive competencies. My results suggest that collaborative ability is more likely to evolve first by between-group conflicts and then later be utilized and improved in games against nature. If collaborative abilities remain low, the species is predicted to become genetically dimorphic with a small proportion of individuals contributing to public goods and the rest free-riding. Evolution of collaborative ability creates conditions for the subsequent evolution of collaborative communication and cultural learning.

## Introduction

Our species is unique in a great variety of different ways but the most crucial of them are related to the size and complexity of our brain (1–6). Brain size in the genus *Homo* tripled in the past 2.5 Myr as a result of several punctuated changes supplemented by gradual within-lineage changes in *Homo erectus* and *Homo sapiens* (2, 7, 8). In modern humans, the brain is very expensive metabolically: it represents about 2% of the body’s weight but utilizes approximately 20% of the energy consumed (8, 9). Other costs include a need for extended parental care due to a longer growth period, difficulties at giving birth to larger-headed babies, and some mental illnesses that come with brain complexity. A burning question is what factors were responsible for the evolution of human brain size and intelligence despite all these costs.

Two sets of explanations have been hotly debated. Ecological explanations include climate variability and harshness, parasites’ and predators’ pressure, as well as changes in diet, habitat use, and food extraction techniques (10, 11). However the empirical support for the role of ecology in human brain evolution is relatively weak. Neocortex size does not seem to correlate with several indices related to diet and habitat (12). There is a statistically significant association of cranial capacity with climate variability and harshness, and parasite pressure, but these factors are much less important than the population density (7).

An alternative set of explanations coming under the rubric of the social brain hypothesis focuses on selective forces resulting from interactions with conspecifics (1, 13). Several types of scenarios have been discussed. One is within-group competition which puts a premium on individuals being able to devise and use “Machiavellian” strategies (including deception, manipulation, alliance formation, exploitation of the expertise of others, etc.) increasing social and reproductive success (6, 14). Comparative studies suggest that species in which Machiavellian-like strategies have been documented have larger brain sizes than related species that do not commonly use these strategies (15). The plausibility of this scenario is also supported by mathematical modeling (16). Another scenario emphasizes selection for the ability to maintain social cohesion in large groups (which become increasingly unstable due to increasing within-group conflicts). It is assumed that larger group sizes are more advantageous because of predatory pressure or in between-group competition. Data do show that brain size correlates with both the group size (12, 17) and population density (7). The third scenario stresses the advantages of social learning over individual learning under conditions of an increasingly fluctuating environment which was characteristic of the Plio-Pleistocene (18). Copying the innovations of others through social learning can be advantageous in such environments especially if the population size is sufficiently high (19). As mentioned above, cranial capacity correlates weakly with environmental variation but strongly with population density (7). Mathematical models do show that the capacity for social learning can increase when an environment changes in spite of its costs (16, 20, 21).

Humans are also unique in their innate ability and willingness to cooperate at a variety of different scales (22, 23). Cooperation often requires efficient collaboration with group-mates which is likely to be very cognitively demanding, especially in conditions that require the rapid and fluid coordination of the behavior of many people, as in hunting or between-group conflict, and the planning for such activities. In fact it has been argued that the evolutionary roots of human cognition are in our capacity to form shared goals, be committed to them, and collaborate in pursuing them and that this capacity evolved within the context of small-group cooperation (23–25) that enhanced competitive ability vis-a-vis that of other groups (3, 5, 26). Within this version of the social intelligence hypothesis, selection for increased ability to collaborate with others (which requires shared goals, joint attention, joint intentions, cooperative communications, etc. (22)) drives the evolution of cognitive abilities. Recent theoretical work has shown that the need for cooperation in dyadic interactions can promote increased brain complexity (27), improved memory (28), and the appearance of tactical deception (29). However because of economies of scale, cooperation and collaboration between multiple social partners can result in significantly larger rewards than that in dyadic interactions, and thus could potentially be a very strong selective force for increased social-cognitive competencies.

There are two general types of collective actions in which our ancestors were almost certainly engaged. The first includes group activities such as defense from predators, some types of hunting or food collection, use of fire, etc. The success of a particular group in these activities largely does not depend on the actions of neighboring groups. I will refer to such collective actions as “us vs. nature” games. The second, which I will refer to as “us vs. them” games, includes conflicts and/or competition with other groups over territory and other resources including mating. The success of a particular group in an “us vs. them” game definitely depends on the actions of other neighboring groups. The outcomes of both types of games strongly affect individual reproduction as well as group survival.

Collective actions often lead to the collective action problem: if individual effort is costly and a group member can benefit from the action of group-mates, then there is an incentive to “free-ride", i.e., reduce one’s effort or withdraw it completely (30–32). But if enough group-mates follow this logic, the public good is not produced and all group members suffer. Overcoming a collective action problem is a major challenge facing many animal and human groups (33–36). During the evolution of our species however, this problem has apparently been solved as humans have strong innate preferences for cooperative and collaborative actions as demonstrated in experiments with small children (25). My goal here is to answer the following questions: Can the need for within-group cooperation and collaboration in collective actions select for increased cognitive abilities overcoming both the collective action problem and various costs of increased intelligence? If yes, which types of collective actions are most conducive in this regard?

A couple of additional clarifications are in order. First, the models to be considered below focus specifically on the evolution of collaborative ability rather than on the evolution of cognitive abilities in general. The former was preceded by a general increase in brain size throughout the Cenozoic in many mammalian lineages (37). Greater brain size is expected to correlate with better cognitive abilities. Second, high cognition obviously has other benefits besides the ability to collaborate. These however are outside of the scope of this paper. Third, my models focus exclusively on social instincts (encoded in genes) and on deeper evolutionary roots of human social behavior. As such, they intentionally neglect the effects of language, culture, and social institutions which are crucial for human ability to cooperate in very large groups.

## Models and Results

I consider a population of individuals living in a large number *G* of groups of constant size *n*. The amount of resources obtained by each group depends on the total effort of its members towards group success; simultaneously individuals pay fitness costs which increase with their efforts. Group members share the reward equally. Generations are discrete and nonoverlapping. A group’s success in solving the collective action problem controls the probability that the group survives to leave offspring to the next generation. The groups that do not survive are replaced by the offspring of surviving groups. Each individual’s effort is controlled genetically and is modeled as a continuous variable. I allow for mutation, recombination, migration, and genetic relatedness between individuals. Below I describe the models and main results. The latter are based on analytical approximations and numerical simulations (see Methods and the Supplementary Information (SI)). My derivations use the assumption that within-population genetic variation is very low (invasion analysis/adaptive dynamics approximation (38, 39)). Numerical simulations show that theoretical conclusions remain valid at a qualitative level even in the presence of genetic variation.

Let *x*_*ij*_ be the effort of individual *i* in group *j* towards the group’s success. For members of surviving groups, I define individual fertility as

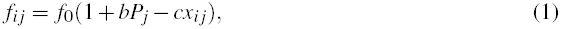

where *b* and *c* are the benefit and cost coefficients, and *f*_0_ is a constant baseline fertility (which can be set to 1 without any loss of generality). As explained below, the probability of the group’s success *P*_*j*_ in producing/securing the public good increases with the group efficiency *X*_*j*_.

A standard practice in evolutionary modeling is to define the group efficiency as an additive function *X*_*j*_ = ∑_*i*_ *x*_*ij*_ of individual efforts. Here I will use a more general and flexible function

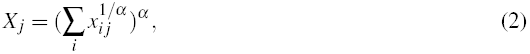

where *α* is a non-negative parameter (40) which I will refer to as collaborative ability.

Collaboration means working with others to achieve shared goals. If individuals are able to collaborate efficiently, a desired outcome can be produced at much smaller individual efforts and/or the same amount of individual effort can result in a much better group outcome than if they acted alone. These intuitions are captured by parameter *α*. In terms of my model, the shared goal is to maximize the group effort *X*_*j*_ which will increase the amount of goods obtained by the group (see below). If collaborative ability *α* is very small (*α* ≪ 1), the group is only as efficient as its member with the largest effort (*X*_*j*_ ≈ max_*i*_(*x*_*ij*_)). Increasing collaborative ability *α* while keeping individual efforts the same increases the group effort *X*_*j*_. If collaborative ability *α* > 1 (*α* < 1), then the group efficiency *X*_*j*_ is larger (smaller) than the sum of individual efforts. If all group members make an equal effort *x*, then *X*_*j*_ = *n*^*α*^ *x*. The latter function is related to the LanchesterOsipov model (41, 42). If individuals vary in their efforts but the variation is relatively small, then 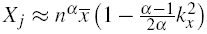, where 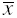 and *k*_*x*_ are the mean and the coefficient of variation of individual efforts *x*. That is, the group efficiency *X*_*j*_ is maximized when the average effort 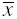 is large but the relative variation in efforts *k*_*x*_ is small, so that there is an increased premium for participation of many individuals.

### Us vs. nature

I start by treating collaborative ability *α* as a constant, exogenously specified parameter. Consider first the “us vs. nature” game. Each group is involved in the production of a public good of value *b* to each member. I define the success probability for group *j* as

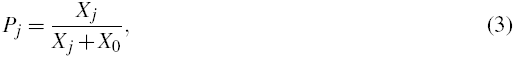

where *X*_0_ is a “half-saturation” parameter (which specifies the group efficiency at which *P*_*j*_ = 50%). The larger *X*_0_, the more group effort *X*_*j*_ is needed to secure the success. I posit that the probability that the group survives to leave the offspring is proportional to its average fertility 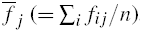. The groups that do not survive are replaced by the offspring of surviving groups. Specifically, I assume that each group in the current generation descends from a group in the previous generation independently with probabilities proportional to 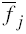. Individuals in each group descend from individuals in their parental group independently with probabilities proportional to *f*_*ij*_. That is, selection is described by a two-level Fisher-Wright framework common in theoretical studies (43). Because each group’s contribution to the next generation depends on its average fitness, this lifecycle corresponds to “hard selection” (44). This is also a model of multilevel selection (45) where group-level selection favors large efforts *x* (which would increase the probability of group success *P*_*j*_) while individual-level selection favors low efforts *x*_*ij*_ (which would reduce the individual costs term *cx*_*ij*_) creating an incentive to free-ride. What makes it beneficial to free-ride is that all group members share equally the benefit of high *P*_*j*_ values independently of their individual contributions to the group’s success.

To understand the model’s behavior it is useful to define the parameter 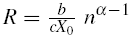. Here the first factor is the ratio of the benefit per individual (*b*) divided by the cost per whole group (*cX*_0_) at a state where the probability of group success *P* = 50%. Under biologically reasonable conditions this ratio will be small. The second factor is an increasing function of *α* which decreases or increases with group size *n* depending on whether *α* < 1 or *α* > 1.

The evolutionary dynamics in this model can be summarized as follows (see SI). If *R* < 1, then at equilibrium individuals make no effort towards the group success (*x*\* = 0). If *R* > 1, then groups make some effort. Specifically, if collaborative ability is low (*α* < *α*_*crit*_), the population is dimorphic with a great majority of individuals making no efforts and a small proportion (approximately one individual per group) making a substantial collaborative effort. If collaborative ability is high (*α* > *α*_crit_), then all individuals will make positive effort. The critical value 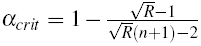 is always smaller than one and approaches one as the group size *n* increases.

The inability of the group to produce the public good when *R* < 1 is a consequence of freeriding exhibited by the group members. In this case, the benefit to cost ratio is too low to secure a positive contribution. If *R* > 1 but collaborative ability *α* is small, the group effort is approximately equal to that of a single individual who is making the largest effort (the “strongest link”). Therefore in this case, for the group to be successful it is sufficient to have a single contributing individual per group. In this model, contributors and free-riders differ genetically. (In contrast, in (46), which also predicted groups composed by a mixture of contributors and free-riders, genetic differences were irrelevant but genes were expressed conditionally depending on exogenous factors.) If *R* > 1 and *α* > *α*_*crit*_, then all individuals start contributing. If group members can effectively collaborate (i.e. *α* > 1), then parameter *R* increases with *n* and the condition *R* > 1 for a positive group effort simplifies. At a state where all group members contribute, the equilibrium individual and group efforts are 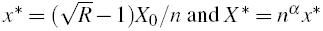. This shows that higher collaborative ability *α* leads both to a higher individual effort *x*\* (via an increase in *R*) and a disproportionately higher group effort *X* * (via the synergistic term *n*^*α*^). Effectively with high *α*, the same benefit can be achieved at smaller individual costs, which removes incentives to free-ride.

### Us vs. them

Next, consider the “us vs. them” game. Assume that groups are involved in direct competition for some resources with total value *bG*. Let the share of the resources obtained by group *j* be (47)

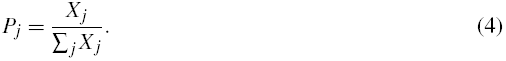

Variable *P*_*j*_ can also be interpreted as a proportion of fights that group *j* won. Losing a conflict can result in group eradication. Assume for simplicity that groups survive to leave offspring to the next generation with probabilities *P*_*j*_.

The behavior of this model is strikingly different from that of the first. Now groups always make a positive effort. If collaborative ability is small (i.e 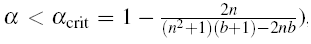, then the group effort is delivered by a small proportion (roughly 1/*n*) of individuals with the rest making no effort. If collaborative ability is large (i.e. *α* > *α*_crit_), then each individual is making effort 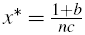. The collaborative ability parameter *α* does not affect the equilibrium level of effort (although the group efficiency *X* * will naturally grow with *α*). These differences between the two models are due to the fact that in the “us vs. nature” games successful production of a public good requires a sufficiently high absolute group effort, while in the “us vs. them” games it is the relative effort that counts, not the absolute. Latter situation creates more favorable conditions for an “arms race” in the individual and group efforts.

### Evolvin *α*

So far I have treated collaborative ability *α* as an exogenously specified constant. Increased ability to collaborate results in a more efficient group effort and, thus, one may expect selection for an increased *α*. To investigate this possibility, assume that each individual is characterized by its own, genetically controlled collaborative ability. I will use the average group collaborative ability in computing the group efficiency *X*_*j*_ (48). To account for individual costs of increased cognitive abilities, I will assume that individual fitness is reduced by a factor 1 − *s*|*α* −θ |, where parameter *s* measures the cost of cognitive abilities and θ is a baseline collaborative ability (which can be positive due to some other benefits of cognitive abilities (3) external to the factors studied here).

To obtain intuition about the strength of forces acting on *α*, assume that the variation in individual efforts *x*_*i*_ is very low. Then in the “us vs. nature” games, analyses show (see SI) that if *x* is very small initially (e.g. maintained by mutation), selection acts against increasing *α*. Selection will act to increase *α* only if individuals make sufficiently large effort in collective action. In contrast, in the “us vs. them” games, the level of individual efforts is irrelevant and there will be selection towards a positive value of *α* (unless costs are very high, specifically *s* > *ln*(*n*)/(*n* − 1)). Collaborative ability is predicted to evolve to 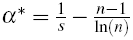. Note that *α*\* does not depend on the costs or benefits of collective action. If the costs are too high, *α* reduces to θ. In both types of games, increasing the group size *n* makes the evolution of collaborative ability more difficult.

The conclusions above are based on simple analytical approximations. To check their validity under more general conditions, I have performed individual-based simulations allowing for the joint evolution of individual efforts and collaborative ability (see SI for details). I assumed that the two traits are controlled by two independent unlinked loci with a continuum of allelic effects. These simulations support my conclusions. In “us vs. nature” games, individuals’ efforts and collaborative abilities increase only if the costs *c* and *s* and the group size *n* are small, benefit *b* is large, and there is a pre-existing high level of collaboration (i.e. θ is high; Fig.1, SI). In “us vs. them” games, collaborative ability and individual efforts evolve under a much broader range of conditions (Fig.2). If collaborative ability does not evolve, then groups become dimorphic as is apparent from increased within-group genetic variation (Fig.2c). Figure 3 illustrates the difference between these two dynamics.

**Figure 1:**
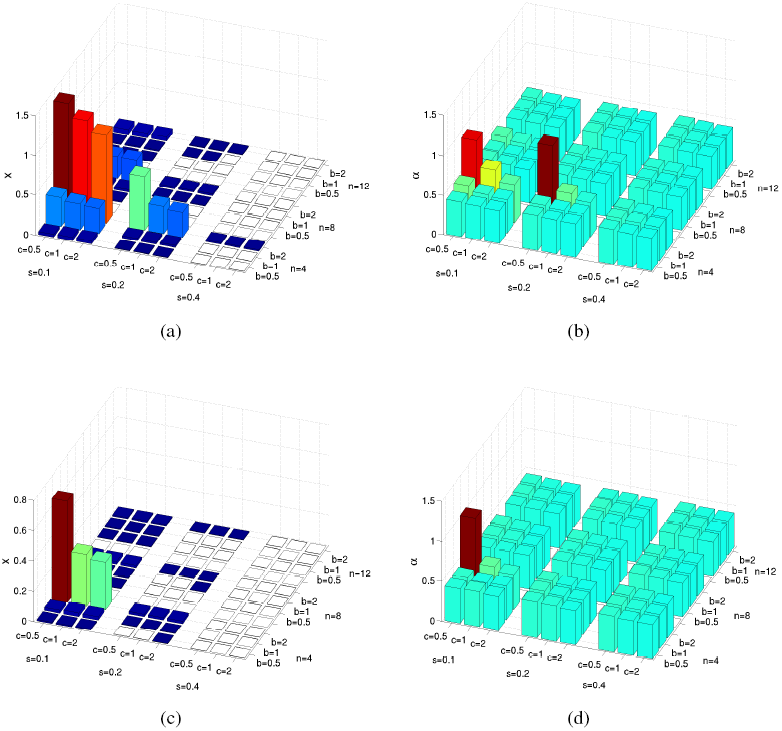
Collective action in “us vs. nature” games. Each figure shows the effects of four parameters (benefit of collaboration *b,* cost of individual effort *c,* cost of collaborative ability *s,* and the group size *n*) on the average equilibrium values of individual effort *x* (first column) and collaborative ability *α* (second column). First row: relatively low individual effort is required for the production of public goods (“half-saturation” parameter *X*_0_ = *0.25n*). Second row: relatively high individual effort is required for the production of public goods (“half-saturation” parameter *X*_0_ = *0.5n*). In most cases, individual effort *x* does not evolve and the collaborative ability *α* remains close to the base-line level of *θ* = 0.4. The height of the bars is also reflected in their color using the jet colormap in *Matlab* (low values in dark blue and high values in brown).

**Figure 2:**
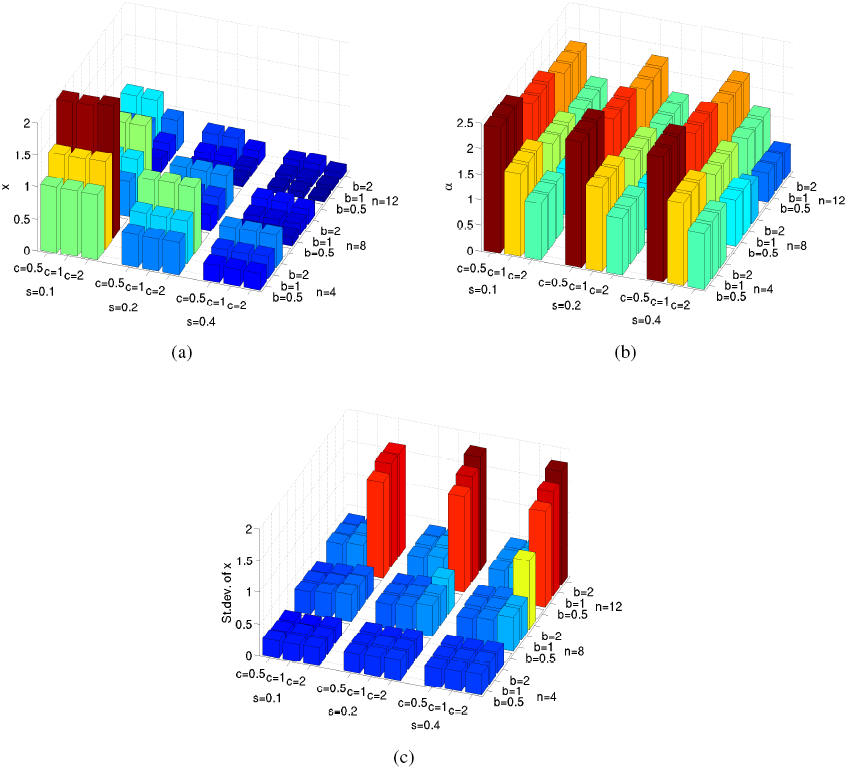
Collective action in “us vs. them” games. Each figure shows the effects of four parameters (benefit of collaboration *b,* cost of individual effort *c,* cost of collaborative ability *s,* and the group size *n*) on (a) Average individual effort at equilibrium *x,* (b) Collaborative ability *α,* and (c) Within-group standard deviation in individual efforts. The base-line collaborative ability *θ* = 0.4. The height of the bars is also reflected in their color using the jet colormap in *Matlab* (low values in dark blue and high values in brown).

**Figure 3:**
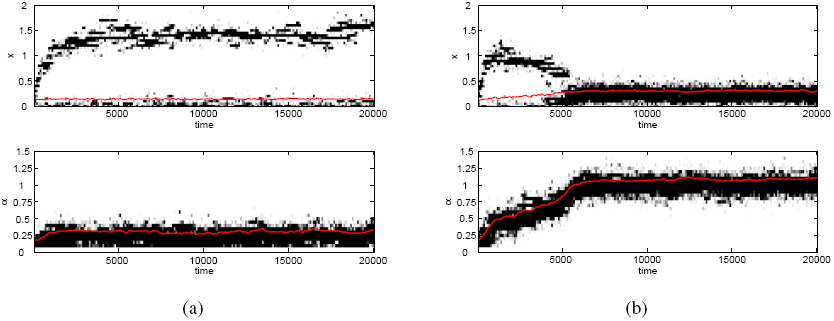
Evolution of individual efforts and collaborative ability in “us vs. them” games. a) High cost of collaborative ability *s* (= 0.4) results in low collaborative ability *α* and bimodal distribution of individual efforts *x*. b) Intermediate cost of collaborative ability *s* (= 0.2) results in the evolution of high collaborative ability *α* and high average efforts *x*. Other parameters: group size *n* = 8, benefit of collaboration *b* = 1, cost of individual effort *c* = 1, and the base-line collaborative ability θ = 0.2. The intensity of the black color is proportional to the number of individuals with the corresponding trait values. Red lines show the mean values.

Overall, my results lead to the following scenario for the evolution of collaborative ability and collective action participation. First, individuals start contributing to collective actions involving direct competition with neighboring groups of conspecifics (“us vs. them” games). Subsequently, they evolve improved ability to collaborate in these actions. Once this ability is established at some level, it becomes used in other collective actions. Specifically, individuals start participating in “us vs. nature” collective games which then produces additional selection for further increases in collaborative ability and social intelligence. Evolution of collaborative ability creates conditions for the subsequent evolution of collaborative communication and cultural learning (25).

## Discussion

Many social organisms living in stable groups often engage in aggressive group interactions with conspecifics from neighboring groups over territory and other resources, including mating opportunities (34, 35, 49–53). Also, many animals often hunt cooperatively in groups (54, 55). In general, even the most complex types of collaboration in animal groups (e.g. those that include division of labor) do not require advanced cognitive abilities and can emerge from very simple behavioral strategies used by individual group members (56, 57). However there are limits on the extent and benefits of simple cooperative acts imposed by the collective action problem. As I show above, evolution of collaborative ability allows groups to mitigate these limits and secure the benefits at much smaller costs.

In the case of cooperative hunting (and other “us vs. nature” games), the theory built here predicts a positive group effort only if both the total cost required to secure the benefit and the group size are relatively small. However, unless the collaborative ability is high, this effort will typically be made by a very small proportion of individuals with the rest contributing almost nothing. Collective effort can potentially evolve if group members have cognitive abilities allowing for efficient collaboration, but metabolic and other costs will typically preclude an increase in cognitive and collaborative abilities. In the case of between-group conflict, groups are predicted to always make a positive effort and typically there will be selection for increased collaborative ability. Only if the cost of cognitive abilities is very high, does collaborative ability not evolve and instead the population becomes dimorphic with a small proportion of individuals making a strong effort towards the group’s success and remaining group members largely free-riding. If high collaborative ability does evolve through between-group conflict, it can be used in “us vs. nature” games which would then further select for increased collaborative ability. This process will lead to a further increase in the effort devoted to “us vs. nature” games and the resulting benefits. In contrast, the effort devoted to between-group conflicts does not depend on the collaborative ability and thus will remain stable as it evolves. Realistically low levels of genetic relatedness will not much affect these conclusions (see SI).

Both types of models considered here include individual and group selection as well as public goods production. However in the “us vs. nature” games, the success of one group in a public goods production does not affect that of another group. In contrast, in “us vs. them” games one group’s success means another group’s failure. This difference results in stronger selection in the “us vs. them” models which in turn produces a larger evolutionary response both in individual contributions and collaborative ability.

The evolution of collaborative ability as studied here requires between-group competition. However strong competition by itself is not enough. In terms of my models, the crucial factors are 1) the level of baseline collaborative ability, 2) benefits and costs of collective actions, 3) costs of collaborative ability, and 4) presence of relevant genetic variation. In the models, collaborative ability evolves only if these four factors are in an adequate range. One can argue then that somehow in the last couple of million years our species found itself under the right conditions. For example, costs of having a big brain might have been reduced as a result of a shift in diet (“expensive tissue” hypothesis, (59)) or there was a shift in the main selective forces from “selection by nature” to “selection by conspecifics” (the “ecological dominance” hypothesis, (1)). My models did not attempt to describe these processes mechanistically but rather captured them in a form of constant parameters.

Starting with Darwin’s *The Descent of Man*, many researchers view between-group conflict and warfare as a potentially important selective factor in shaping many human characteristics (Ref.(60–62) but see Ref.(63)). In particular, it has been argued that between-group conflict was a driving force in the emergence of many human abilities, biases, and preferences (such as cooperation, belligerence, leadership, altruism, parochialism, and ethnocentrism) as well as human social norms and institutions (64–67, 46). Alexander (26) argued that the need to succeed in between-group competition would select for increased human cognition and mental abilities; thus allowing for more concerted and effective group actions. Here, I have provided strong theoretical support to these arguments by showing that between-group conflict can select for increased intelligence and cognitive abilities used to coordinate group activities, potentially overcoming both the high costs of large brains and the collective action problem.

It is also generally believed that large-game hunting was very important in human evolution. Success in large-game hunting required the consistent coordinated collaboration of multiple hunters. Alexander (26) argued that collaboration in hunting came first and subsequently created conditions for the evolution of collaboration in between-group conflicts (see also Ref.(25)). My results however show that the reverse sequence (i.e. collaboration in between-group fighting followed by that in large-game hunting (3)) is more plausible. Collaboration and commitment to a shared goal are also very important in within-group coalitions and alliances which represent an efficient form of within-group competition for reproductive success in a number of mammals including hyenas, wolves, dogs, lions, cheetahs, coatis, meerkats, various primates and dolphins (68–70). My results suggest that within-group coalitions were preceded and promoted by between-group conflicts. Both these hypotheses still require empirical substantiation. Also it has been argued that human cognitive evolution was driven by selection for cooperative breeding (71). The latter scenario largely relies on indirect (72) rather than on direct benefits as considered here and therefore is less likely. However, once collaborative ability and shared intentionality are established in the species, the evolution of cooperative breeding is greatly simplified.

The prediction of within-group polymorphism in animals with low collaborative ability and no shared intentionality is supported by an observation that most effort in chimpanzees’ group activities is provided by a small number of “impact hunters” and “impact patrollers” (73). The prediction that humans have evolved a genetic predisposition for collaborative group activities is in line with a consistent observation that human infants are motivated to collaborate in pursuing a common goal (22) and that cooperative acts result in activation of brain regions involved in reward processing, independently of material gains (74). People cooperate when groups face failure due to external threats, e.g. harsh environmental conditions or natural disasters (75, 76). However, as predicted by the theory above, cooperation increases dramatically in the presence of direct between-group competition (77–82) to a level that “cues of group competition have an automatic or unconscious effect on human behaviour that can induce increased within-group cooperation” (81). A variety of other facts and observations about human psychology (e.g. in-group/out-group biases, widespread obsession with team sports, and sex differences in the motivation to form and skill at maintaining large competitive groups (3)) strongly support the idea about the importance of between-group conflicts in shaping human social instincts.

The models presented here can be extended and generalized in a number of ways. For example, it is known that the outcome of multilevel selection can depend on the frequency of group reproduction events. In the models presented, group reproduction happens every generation according to the Fisher-Wright scheme. An open question is to what extent the results hold up when group reproduction happens less frequently. Also, one can use alternative functions to represent the strength of the group (eq.2) and the costs of collaborative ability, or introduce variation between groups members in, say, their strength or valuation of the reward (46), or assume that individual cognitive abilities affect the costs they pay, etc. Future studies of these and similar modification are needed to increase the generality of this approach and shed additional light on the evolution of human cooperation and cognition.

Decades of intensive work by generations of evolutionary biologists have led to a dramatic increase of our knowledge of how new species arise (83, 84). Time is ripe for a systematic effort to understand the ultimate speciation process that of our own species (85). Identifying evolutionary roots for and the dynamics of both human cognitive abilities and cooperative social instincts is a necessary step in getting a better understanding of the origins of our “uniquely unique” species (1).

## Methods

In numerical simulations, all individuals were sexual haploids; each deme comprised *n* males and *n* females. Only males contributed to the public good production and paid individual costs. Females carried the genes for the amount of effort and collaborative ability but they were not expressed. Each group in the current generation descended from a group in the previous generation randomly and independently with probability 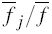 (in the “us vs. nature” game) or *P*_*j*_ (in the “us vs. them” game). To populate a “descending” group, each female in the corresponding “ancestral” group produced two offspring. The fathers where chosen randomly and independently from the pool of the group’s males with probabilities 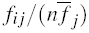. I assumed free recombination between the two genes. The offspring sex was assigned randomly but within each group I enforced an equal sex ratio. Female offspring dispersed randomly between demes while male offspring stayed in the native deme. Simulations ran for 200,000 generations.

In numerical studies of the basic model I used all possible combinations of the following parameter values: expected benefit per individual *b* = 0.5, 1.0, 2.0; cost coefficient *c* = 0.5, 1.0, 2.0; group size *n* = 4, 8, 12; base-line collaborative ability θ = 0.1, 0.2, 0.4, 0.8, and in “us vs. nature” model half-effort parameter *x*_0_ = 0.25, 0.5, 1.0, 2.0. I performed 10 runs for each parameter combination. Some parameters did not change: number of groups *G* = 1000, mutation rate *μ* = 0.001 per gene per generation, standard deviation of the mutational effect *σ*_*μ*_ = 0.1. The initial individual efforts were chosen randomly and independently from a uniform distribution on [0, 0.05]. The initial value of collaborative ability were chosen randomly and independently from a uniform distribution on [θ, θ + 0.05]. To avoid the appearance of negative fitness values in numerical simulations, I introduced upper boundary on individual efforts *x*_max_ = (1 + *b*)/*c*. I used zero lower boundary on *x*. Supplementary Figures 1-5 summarize the results.

## Acknowledgments

I thank E. Akcay, P. Barclay, L. Fortunato, D. C. Geary, K. Rooker, and reviewers for comments, suggestions, and discussions. Supported in part by the National Institute for Mathematical and Biological Synthesis through NSF Award #EF-0830858, with additional support from The University of Tennessee, Knoxville, and by the U.S. Army Research Laboratory and the U. S. Army Research Office under grant number W911NF-14-1-0637.

## Supplementary Information for S. Gavrilets “Collective action and the collaborative brain”

To study my models I used the evolutionary invasion analysis (adaptive dynamics (1, 2)) which focuses on the invasion fitness *w*(*y*|*x*), i.e., the fitness of a rare mutant with trait *y* in a monomorphic population of individuals with trait *x*. In the equations describing my main results, I will assume that the number of groups *G* is large.

### Games against nature

Under the assumptions stated in the main text, fitness of individual *i* from group *j* is

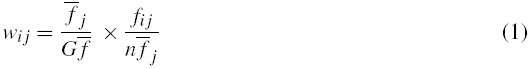

where the first and the second ratios are the probability of group survival and the share of the group reproduction going to the individual, respectively, with 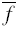 being the average group fertility in the population (i.e. 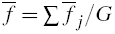).

#### Additivity

Assume that the group efficiency function is additive, i.e. *X*_*j*_ = ∑_*i*_*x*_*ij*_. Then invasion fitness can be written as:

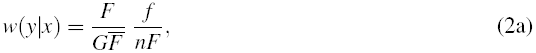

where the mutant’s fertility *f*, the average fertility of the mutant’s group *F*, and the average fertility of groups in the population 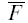). Are

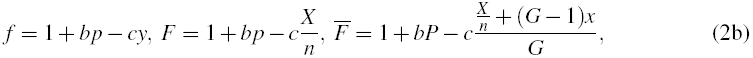

with

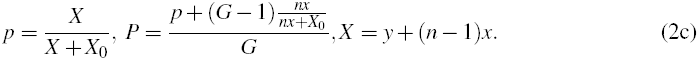

Here, *p* and *P* are the probability of success for the mutant’s group and the average probability of success in the population, respectively, and *X* is the efficiency of the mutant’s group.

Making a variable change *z* = *x*/*x*_0_ with *x*_0_ = *X*_0_/*n*, we find that the selection gradient

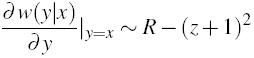

where 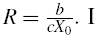. I conclude that if *R* > 1, the individual effort evolves to an equilibrium value

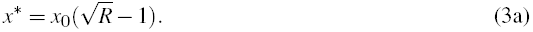

This equilibrium is stable. If *R* < 1, then *x*\* = 0.

#### Synergicity

Assume that the group efficiency function is

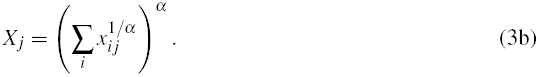

This assumption results in the equations for *P* and *X* becoming

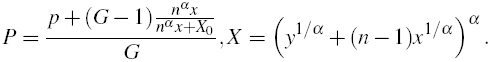

Using the same approach as above, one finds that the equation for *R* becomes

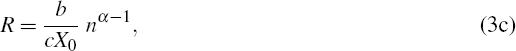

while the nonzero equilibrium value of *x* is still given by expression (3a). However this equilibrium is stable only if

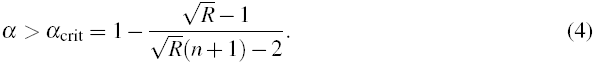

If the above condition is not satisfied, *x*\* is a branching point (1, 2) and the population becomes dimorphic with most individuals contributing nothing and on average one individual per group making a large nonzero contribution.

#### Evolution of *α*

To study the effects of selective forces acting on collaborative ability *α*, we assume that individuals are monomorphic with respect to their effort *x*. I also assume that viability decreases linearly with collaborative ability. Then the fitness of a rare mutant with collaborative ability *β* in a resident population with ability *α* can be written as:

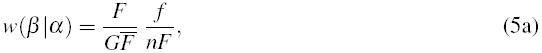

where the mutant’s fertility *f*, the average fertility of the mutant’s group *F*, and the average fertility of groups in the population 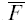 are

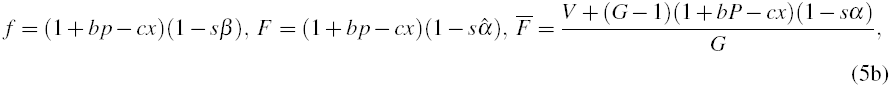

with

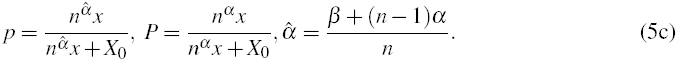

Here, *p* and *P* are the probability of success for the mutant’s group and for a group with no mutant, respectively, and 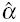 is the collaborative ability of the mutant’s group.

Computing the selection gradient 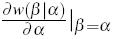 we find that if individual effort *x* = 0, then the selection gradient is 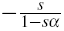 so that *α* will decrease to zero. This suggests that collaboration effort will experience positive selection only if *x* is sufficiently large.

### Games against other groups

Under the assumptions stated in the main text, fitness of individual *i* from group *j* is

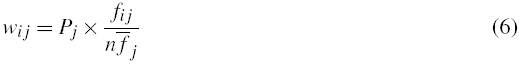

#### Additivity

Assume that the group efficiency function is additive, i.e. *X*_*j*_ = ∑_*i*_*x*_*ij*_. Then invasion fitness (i.e. fitness of a rare mutant with trait *y* in a resident population with trait *x*) can be written as:

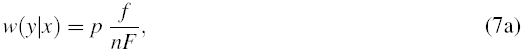

where the mutant’s fertility *f*, the average fertility of the mutant’s group *F*, and the probability of success *p* for the mutant’s group are

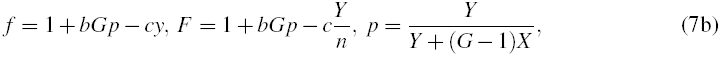

with

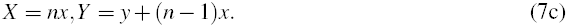

Here, *X* and *Y* are the group efficiencies for the resident and mutant groups, respectively.

Computing the selection gradient, one finds that *x* evolves towards a stable equilibrium which, in the limit of large *G*, can be written as

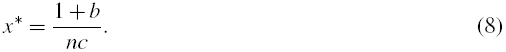

#### Synergicity

Assume that the group efficiency function is given by equation (3b). Then the expressions for *X* and *Y* above take form

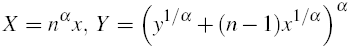

Computing the selection gradient I find that there is still an equilibrium at *x*\* = (1 + *b*)/(*nc*). However, this equilibrium is stable only for *α* large than

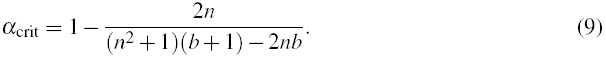

Note that *α*_crit_ approaches zero from below as *n* and/or *b* increase. For *α* < *α*_crit_, *x*\* is a branching point (1, 2). Numerical simulations show that in these case, the population becomes dimorphic, with most individuals contributing nothing and on average 1 individual per group making a large nonzero contribution.

#### Evolution of *α*

Assume first that the population is monomorphic with respect to *x*. The invasion fitness of a rare mutant with collaborative ability *β* in a resident population with ability *α* can be written as:

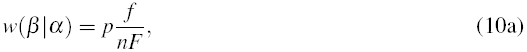

where the mutant’s fertility *f* and the average fertility of the mutant’s group *F* are

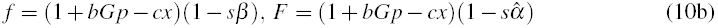

with

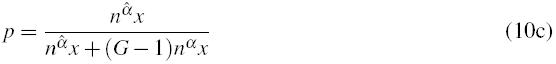

being the probability of the mutant’s group survival. Computing the selection gradient, one finds that the evolutionary dynamics of the collaborative ability *α* do not depend on the individual effort *x*, and that the collaborative ability is predicted to evolve to

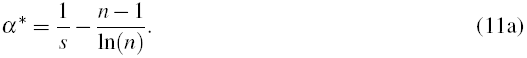

This value is positive if

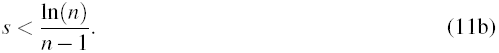

Assume next that the population is dimorphic so that on average only *m* out of *n* group members make effort *x* while the remaining group members make no effort.The equations for the invasition fitness stay the same except that probability *p* of the mutant group success becomes

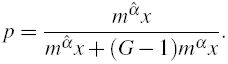

I find that the equilibrium value of *α* becomes

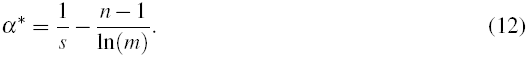

This value is positive if

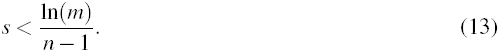

Thus, decreasing the number *m* of contributing group members, makes the evolution of collaborative ability more difficult.

### Relatedness

The results above assume that the groups are formed randomly each generation, implying that group members are genetically unrelated. Evolution of cooperation in public goods games studied here is driven by overlapping interests and does not require genetic relatedness (or reciprocity or punishment). Genetic relatedness does however increase cooperation (3, 4). For situations when all individuals contribute in the “us vs. nature” game, the composite parameter *R* is increased by a factor of 1 + *r*(*n* − 1), where *r* is the average within-group relatedness. This will increase both the range of parameter values resulting in nonzero group efforts and that effort itself. In the “us vs. them” game, each individual effort increases by a factor of 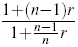. For example, let female offspring disperse randomly between groups while the male offspring stay in their native group (as in chimpanzees and, likely, our ancestors (5, 6)). Then males within a group will be genetically related with 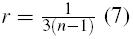 (7). This will result in about a 30% increase in all individual efforts (assuming that *n* ≥ 5).

### Details of numerical simulations

All individuals were sexual haploids; each deme comprised *n* males and *n* females. Only males contributed to the public good production and paid individual costs. Females carried the genes for the amount of effort and collaborative ability but they were not expressed. Each group in the current generation descended from a group in the previous generation randomly and independently with probability 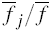 (in the “us vs. nature” game) or *P*_*j*_ (in the “us vs. them” game). To populate a “descending” group, each female in the corresponding “ancestral” group produced two offspring. The fathers where chosen randomly and independently from the pool of the group’s males with probabilities 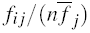. I assumed free recombination between the two genes. The offspring sex was assigned randomly but within each group I enforced an equal sex ratio. Female offspring dispersed randomly between demes while male offspring stayed in the native deme. Simulations ran for 200,000 generations.

In numerical studies of the basic model I used all possible combinations of the following parameter values: expected benefit per individual *b* = 0.5, 1.0, 2.0; cost coefficient *c* = 0.5, 1.0, 2.0; group size *n* = 4, 8, 12; base-line collaborative ability θ = 0.1, 0.2, 0.4, 0.8, and in “us vs. nature” model half-effort parameter *x*_0_ = 0.25, 0.5, 1.0, 2.0. I performed 10 runs for each parameter combination. Some parameters did not change: number of groups *G* = 1000, mutation rate *μ* = 0.001 per gene per generation, standard deviation of the mutational effect *σ*_*μ*_ = 0.1. The initial individual efforts were chosen randomly and independently from a uniform distribution on [0, 0.05]. The initial value of collaborative ability were chosen randomly and independently from a uniform distribution on [θ, θ + 0.05]. To avoid the appearance of negative fitness values in numerical simulations, I introduced upper boundary on individual efforts *x*_max_ = (1 + *b*)/*c*. I used zero lower boundary on *x*. Figures S1–S5 summarize the results.

**Figure 1:**
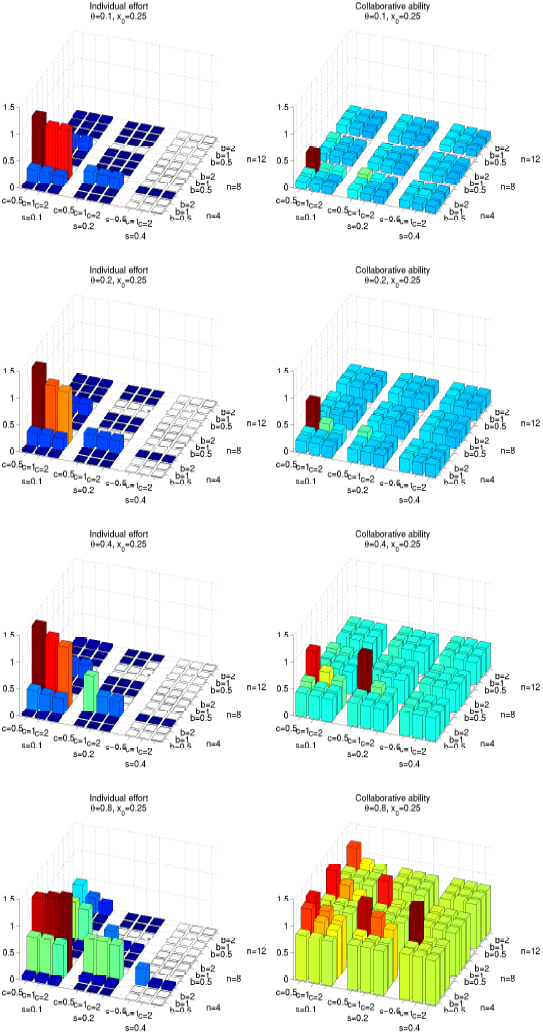
Collective action in “us vs. nature “ games with *x*_0_ = 0.25.

**Figure 2:**
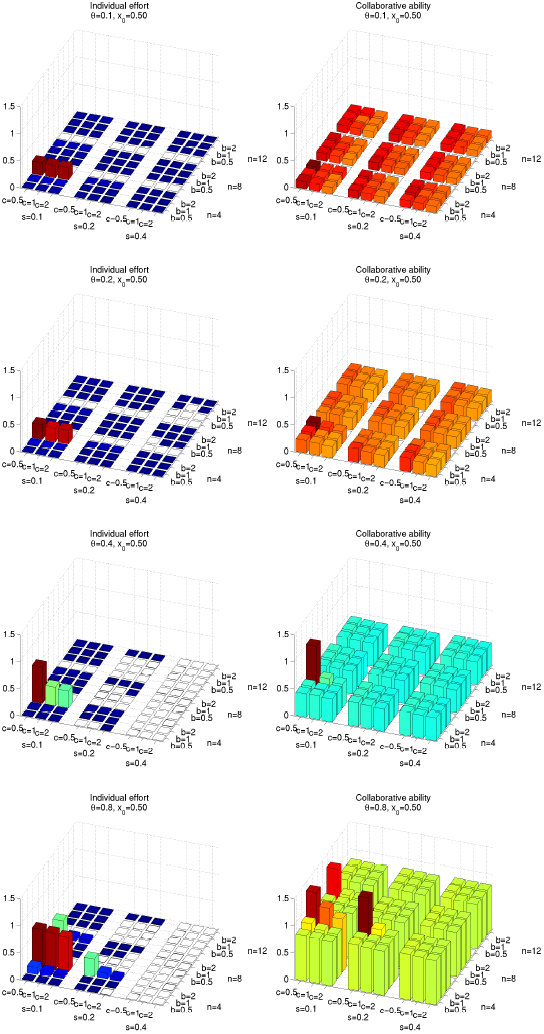
Collective action in “us vs. nature” games with *x*_0_ = 0.5.

**Figure 3:**
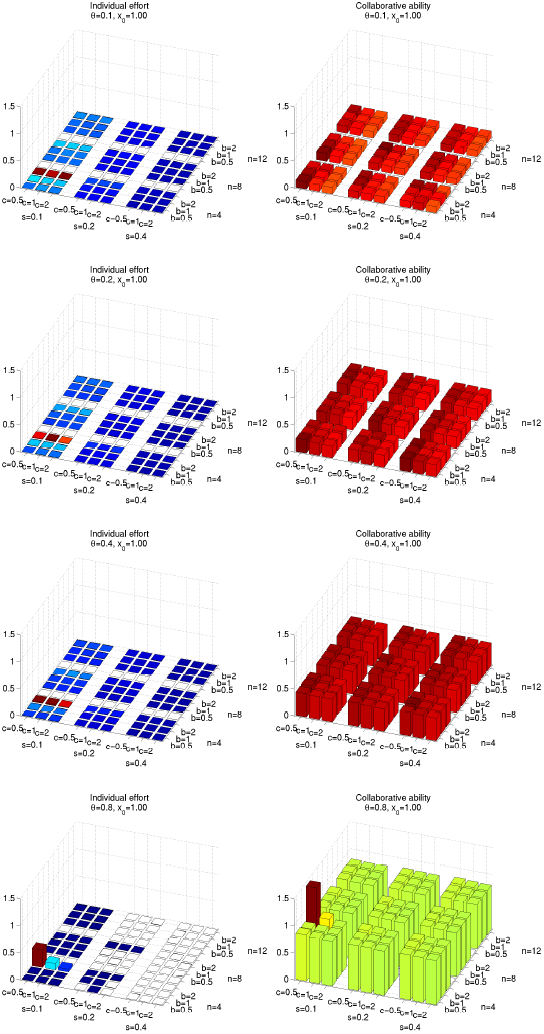
Collective action in “us vs. nature” games with *x*_0_ = 1.

**Figure 4:**
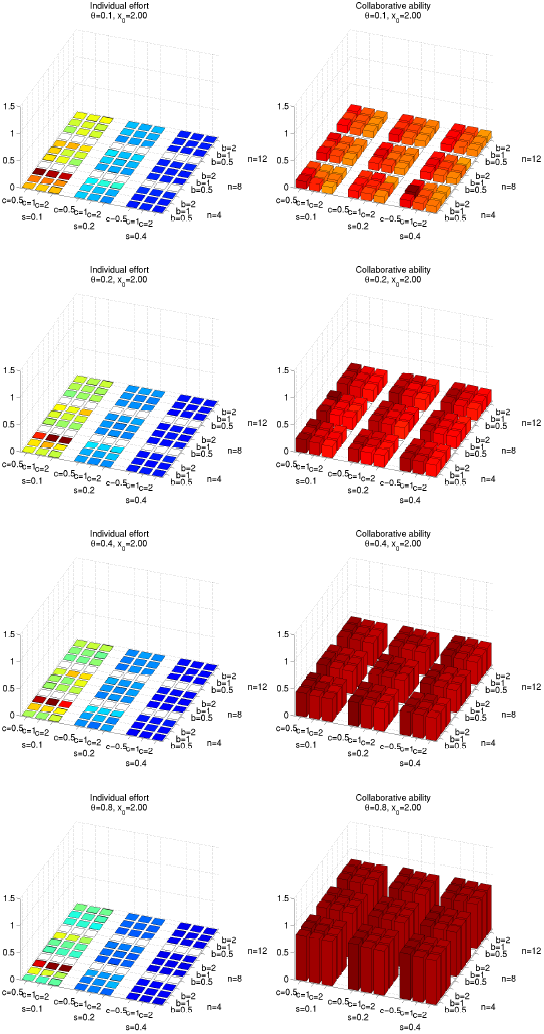
Collective action in “us vs. nature” games with *x*_0_ = 2.

**Figure 5:**
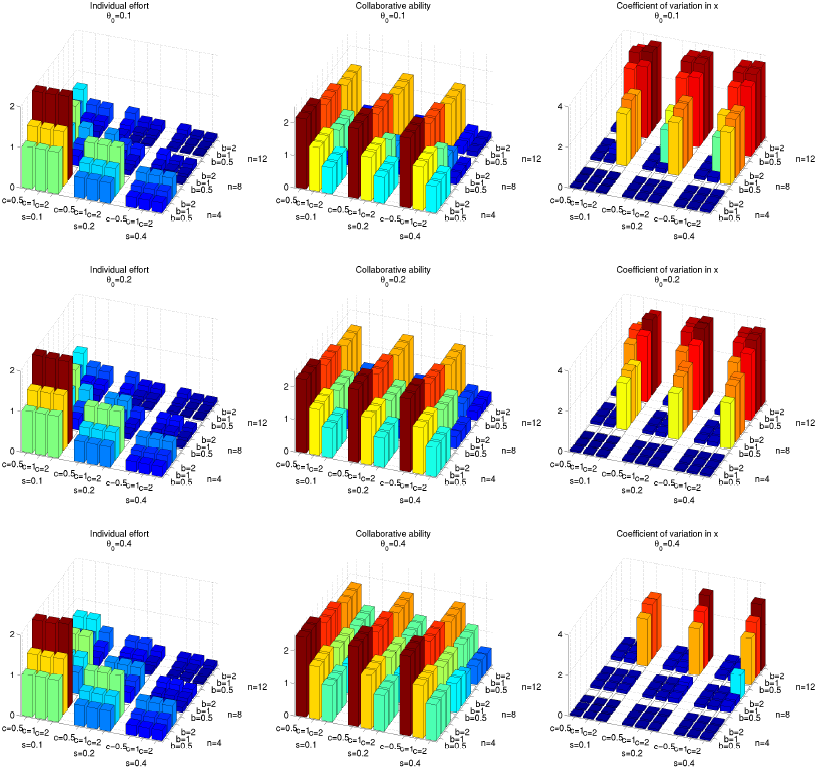
Collective action in “us vs. them” games.

## References

[1] Alexander, R. D. How did humans evolve? Reflections on the uniquely unique species (University of Michigan, Museum of Zoology, 1990).

[2] Striedter, G. F. Principles of brain evolution (Sinauer, Sunderland, MA, 2005).

[3] Geary, D. C. The origin of mind. Evolution of brain, cognition, and general intelligence (American Psychological Association, Washington, DC, 2010).

[4] Roth, G. & Dicke, U. Evolution of the brain and intelligence. Trends in Cognitive Sciences 9, 250–257 (2005).

[5] Flinn, M. V., Geary, D. C. & Ward, C. V. Ecological dominance, social competition, and coalitionary arms races: why humans evolved extraordinary intelligence? Evolution and Human Behavior 26, 10–46 (2005).

[6] Whiten, A. & Erdal, D. The human socio-cognitive niche and its evolutionary origins. Philosophical Transactions of the Royal Society London B 367, 2119–2129 (2011).

[7] Bailey, D. H. & Geary, D. C. Hominin brain evolution: Testing climatic, ecological, and social competition models. Human Nature 20, 67–79 (2009).

[8] Shultz, S., Nelson, E. & Dunbar, R. I. M. Hominin cognitive evolution: identifying patterns and processes in the fossil and archaeological record. Philosophical Transactions of the Royal Society London B 367, 2130–2140 (2012).

[9] Holloway, R. Evolution of the human brain. In Lock, A. & Peters, C. R. (eds.) Handbook of human symbolic evolution, 74–125 (Clarendom Press, Oxford, 1996).

[10] Vrba, E. S. The fossil record of African antelopes (Mammalia, Bovidae) in relation to human evolution and paleoclimate. In Vrba, E. S., Denton, G. H., Partridge, T. C. & Burckle, L. H. (eds.) Paleoclimate and evolution, with emphasis on human origins, 385–424 (Yale University Press, New Haven, CT, 1995).

[11] Russon, A. E. & Begun, D. R. The evolution of thought. Evolutionary origins of great ape intelligence (Cambridge University Press, Cambridge, 2004).

[12] Dunbar, R. I. M. The social brain hypothesis. Evolutionary Anthropology 6, 178–190 (1998).

[13] Holloway, R. L. The role of human social behavior in the evolution of the brain (James Arthur lecture on the evolution of the human brain, no. 43 (American Museum of Natural History, New York, 1973).

[14] Whiten, A. & Byrne, R. W. Machiavellian intelligence II. Extensions and evaluations (Cambridge University Press, Cambridge, 1997).

[15] Byrne, R. W. & Corp, N. Neocortex size predicts deception rate in primates. Proceedings of the Royal Society London B 271, 1693–1699 (2004).

[16] Gavrilets, S. & Vose, A. The dynamics of Machiavellian intelligence. Proceedings of the National Academy of Sciences USA 103, 16823–16828 (2006).

[17] Dunbar, R. I. M. The social brain hypothesis and its implications for social evolution. Annals of Human Biology 36, 562–572 (2009).

[18] Richerson, P. J., Bettinger, R. L. & Boyd, R. Evolution on a restless planet: were environmental variability and environmental change major drivers of human evolution. In Woketits, F. M. & Ayala, F. J. (eds.) Handbook of evolution. Vol. 2. The evolution of living systems, 223–242 (Wiley, 2005).

[19] Richerson, P. J. & Boyd, R. Not By Genes Alone. How Culture Transformed Human Evolution (University of Chicago Press, Chicago, 2005).

[20] Higgs, P. G. The mimetic transition: a simulation study of the evolution of learning by imitation. Proceedings of the Royal Society London B 267, 1355–1361 (2000).

[21] Nakahashi, W. Evolution of learning capacities and learning levels. Theoretical Population Biology 78, 211–224 (2010).

[22] Moll, H. & Tomasello, M. Cooperation and human cognition: the Vygotskian intelligence hypothesis. Philosophical Transactions of the Royal Society London B 362, 639–648 (2007).

[23] Tomasello, M. & Moll, H. The Gap is Social: Human Shared Intentionality and Culture. In Kappeler, PM and Silk, JB (ed.) Mind the gap: tracing the origins of human universals, 331–349 (Springer-Verlag, 2010).

[24] Melis, A. P. & Semmann, D. How is human cooperation different. Philosophical Transactions of the Royal Society London B 365, 2663–2674 (2010).

[25] Tomasello, M., Melis, A. P., Tennie, C., Wyman, E. & Herrmann, E. Two key steps in the evolution of human cooperation. Current Anthropology 53, 673–692 (2012).

[26] Alexander, R. D. Evolution of the human psyche. In Mellars, P. & Stringer, C. (eds.) The human revolution. Behavioural and biological perspectives on the origin of modern humans, 455–513 (Princeton University Press, Princeton, NJ, 1989).

[27] McNally, L., Brown, S. P. & Jackson, A. L. Cooperation and the evolution of intelligence. Proceedings of the Royal Society London B 1740, 3027–3034 (2012).

[28] Moreira, J. et al. Individual memory and the emergence of cooperation. Animal Behaviour 85, 233–239 (2013).

[29] McNally, L. & Jackson, A. L. Cooperation creates selection for tactical deception. Proceedings of the Royal Society London B 280 (2013).

[30] Olson, M. Logic of collective action: Public goods and the theory of groups (Harvard University Press, Cambridge, MA, 1965).

[31] Ostrom, E. Governing the commons. The evolution of institutions for collective action (Cambridge University Press, Cambridge, 1990).

[32] Batina, R. G. & Ihori, T. Public goods. Theories and evidence (Solar Phys., Berlin, 2005).

[33] Nunn, C. Collective action, free-riding, and male extra-group conflict. In Kappeler, P. (ed.) Primate males, 192–204 (Cambridge University Press, Cambridge, 2000).

[34] Kitchen, D. M. & Beehner, J. C. Factors affecting individual participation in group-level aggression among non-human primates. Behaviour 144, 1551–1581 (2007).

[35] Willems, E., Hellriegel, B. & van Schaik, C. P. The collective action problem in primate territory conflicts. Proceedings of the Royal Society London B 280 (2013).

[36] Carballo, D. M. (ed.) Cooperation and collective action. Archaeological perspectives. (University Press of Colorado, Boulder, 2013).

[37] Jerison, H. J. Evolution of the Brain and Intelligence (Academic Press, New York, 1973).

[38] Geritz, S. A. H., Kisdi, E., Meszéna, G. & Metz, J. A. J. Evolutionary singular strategies and the adaptive growth and branching of the evolutionary tree. Evolutionary Ecology 12, 35–57 (1998).

[39] Waxman, D. & Gavrilets, S. Target review: 20 questions on adaptive dynamics. Journal of Evolutionary Biology 18, 1139–1154 (2005).

[40] Hirshleifer, J. The paradox of power. Economics & Politics 3, 177–200 (1991).

[41] Gavrilets, S., Duenez-Guzman, E. A. & Vose, M. D. Dynamics of coalition formation and the egalitarian revolution. PLOS One 3, e3293 (2008).

[42] Gavrilets, S. On the evolutionary origins of the egalitarian syndrome. Proceedings of the National Academy of Sciences USA 109, 14069–14074 (2012).

[43] Schonmann, R. H., Vicente, R. & Caticha, N. Altruism can proliferate through population viscosity despite high random gene flow. PLOS One 8 (2013).

[44] Whitlock, M. Selection and drift in metapopulations. In Hanski, I. & Gaggiotti, O. (eds.) Ecology, Genetics, and Evolution in Metapopulations, 153174 (Elsevier, Amsterdam, 2004).

[45] Okasha, S. Evolution and the Levels of Selection (Oxford University Press, Oxford, 2009).

[46] Gavrilets, S. & Fortunato, L. Evolution of social instincts in between-group conflict with within-group inequality. Nature Communications 5, article 3526 (doi:10.1038/ncomms4526) (2014).

[47] Konrad, K. Strategy and dynamics in contests (Oxford University Press, Oxford, 2009).

[48] Devine, D. & Philips, J. Do smarter teams do better - A meta-analysis of cognitive ability and team performance. Small Group Research 32, 507–532 (2001).

[49] Watts, D. P. & Mitani, J. C. Boundary patrols and intergroup encounters in wild chimpanzees. Behaviour 138, 299–327 (2001).

[50] Mosser, A. & Packer, C. Group territoriality and the benefits of sociality in the Athrican lion, Panthera leo. Animal Behaviour 78, 359–370 (2009).

[51] Mitani, J. C., Watts, D. P. & Amsler, S. J. Lethal intergroup aggression leads to territorial expansion in wild chimpanzees. Current Biology 20, R507–R508 (2010).

[52] Mares, R., Young, A. & Clutton-Brock, T. Individual contributions to territory defense in a cooperative breeder: weighing up the benefits and the costs. Proceedings of the Royal Society London B 279, 3989–3995 (2013).

[53] Wilson, M. L., Kahlenberg, S. M., Wells, M. & Wrangham, R. W. Ecological and social factors affect the occurrence and outcomes of intergroup encounters in chimpanzees. Animal Behavior 83, 277–291 (2012).

[54] Packer, C. & Ruttan, L. Evolution of cooperative hunting. American Naturalist 132, 159–198 (1988).

[55] Boesch, C. Cooperative hunting in wild chimpanzees. Animal Behavior 48, 653–667 (1994).

[56] Bailey, I., Myatt, J. P. & Wilson, A. M. Group hunting within the Carnivora: physiological, cognitive and environmental influences on strategy and cooperation. Behavioral Ecology and Sociobiology 67, 1–17 (2013).

[57] Brosnan, S. F., Salwiczek, L. & Bshary, R. The interplay of cognition and cooperation. Philosophical Transactions of the Royal Society London B 365, 2699–2710 (2010).

[58] Tomasello, M., Carpenter, M., Call, J., Behne, T. & Moll, H. Understanding and sharing intentions: The origins of cultural cognition. Behavioral and Brain Sciences 28, 675+ (2005).

[59] Aiello, L. C. & Wheeler, P. The expensive-tissue hypothesis. The brain and the digestive system in human and primate evolution. Current Anthropology 36, 199–221 (1995).

[60] Keeley, L. War before civilization (Oxford University Press, New York, 1996).

[61] Bowles, S. Did warfare among ancestral hunter-gatherers affect the evolution of human social behaviors? Science 324, 1293–1298 (2009).

[62] Gat, A. War in human civilization (Oxford University Press, Oxford, 2006).

[63] Culotta, E. Latest skirmish over ancestral violence strikes blow for peace. Science 341, 224 (2013).

[64] Bowles, S. & Gintis, H. A Cooperative Species: Human Reciprocity and Its Evolution (Princeton University Press, 2011).

[65] Lehmann, L. & Feldman, M. W. War and the evolution of belligerence and bravery. Proceedings of the Royal Society London B 275, 2877–2885 (2008).

[66] Gavrilets, S., Anderson, D. G. & Turchin, P. Cycling in the complexity of early societies. Cliodynamics: The Journal of Theoretical and Mathematical History 1, 5536t55r (2010).

[67] Turchin, P., Currie, T., Turner, E. & Gavrilets, S. War, space, and the evolution of Old World complex societies. Proceedings of the National Academy of Sciences USA 109, 14069–14074 (2013).

[68] Harcourt, A. H. & de Waal, F. B. M. Coalitions and alliances in humans and other animals (Oxford University Press, Oxford, 1992).

[69] Olson, L. & Blumstein, D. A trait-based approach to understand the evolution of complex coalitions in male mammals. Behav Ecol 20, 624–632 (2009).

[70] Smith, J. et al. Evolutionary forces favoring intragroup coalitions among spotted hyaenas and other animals. Behavioral Ecology 21, 284–303 (2010).

[71] Burkart, J. M., Hrdy, S. B. & van Schaik, C. Cooperative breeding and human cognitive evolution. Evolutionary Anthropology 18, 175–186 (2009).

[72] Lukas, D. & Clutton-Brock, T. Cooperative breeding and monogamy in mammalian societies. Proceedings of the Royal Society London B 279, 2151–2156 (2012).

[73] Gilby, I. C., Wilson, M. L. & Pusey, A. E. Ecology rather than psychology explains co-occurrence of predation and border part ols in male chimpanzees. Animal Behaviour 86, 61–74 (2013).

[74] Rilling, J. K. The Neurobiology of Cooperation and Altruism. In Sussman, RW and Cloninger, CR (ed.) Origins of Altruism and Cooperation, Developments in Primatology-Progress and Prospects, 295–306 (Springer-Verlag, 2011).

[75] Vasi, I. & Macy, M. The mobilizer’s dilemma: Crisis, empowerment, and collective action. Social Forces 81, 979–998 (2003).

[76] Milinski, M., Sommerfeld, R. D., Krambeck, H.-J., Reed, F. A. & Marotzke, J. The collective-risk social dilemma and the prevention of simulated dangerous climate change. Proceedings of the National Academy of Sciences USA 105, 2291–2294 (2008).

[77] Gunnthorsdottira, A. & Rapoport, A. Embedding social dilemmas in intergroup competition reduces free-riding. Organizational Behavior and Human Decision Processes 101, 184–199 (2006).

[78] Tan, J. H. W. & Bolle, F. Team competition and public goods game. Economics Letters 96, 133–139 (2007).

[79] Puurtinen, M. & Mappes, T. Between-group competition and human cooperation. Proceedings of the Royal Society London B 276, 355–360 (2009).

[80] Burton-Chellew, M. N., Ross-Gillespie, A. & West, S. A. Cooperation in humans: competition between groups and proximate emotions. Evolution and Human Behavior 31, 104–108 (2010).

[81] Burton-Chellew, M. N. & West, S. A. Pseudocompetition among groups increases human cooperation in a public-goods game. Animal Behavior 84, 947–952 (2012).

[82] Egas, M., Kats, R., van der Sar, X., Reuben, E. & Sabelis, M. W. Human cooperation by lethal group competition. Scientific Reports 3:1373, 1–3 (2013).

[83] Coyne, J. & Orr, H. A. Speciation (Sinauer Associates, Inc., Sunderland, Massachusetts, 2004).

[84] Gavrilets, S. Fitness landscapes and the origin of species (Princeton University Press, Princeton, NJ, 2004).

[85] Darwin, C. The Descent of Man, and Selection in Relation to Sex (John Murray, London, 1871).

## References

[1] Geritz, S. A. H., Kisdi, E., Meszéna, G. & Metz, J. A. J. Evolutionary singular strategies and the adaptive growth and branching of the evolutionary tree. Evolutionary Ecology 12, 35–57 (1998).

[2] Waxman, D. & Gavrilets, S. Target review: 20 questions on adaptive dynamics. Journal of Evolutionary Biology 18, 1139–1154 (2005).

[3] Nowak, M. Evolutionary dynamics (Harvard University Press, Harvard, 2006).

[4] McElreath, R. & Boyd, R. Mathematical models of social evolution. A guide for the perplexed (Chicago University Press, Chicago, 2007).

[5] Copeland, S. R. et al Strontium isotope evidence for landscape use by early hominins. Nature 474, 7678 (2011).

[6] Laluenza-Fox, C. et al Genetic evidence for patrilocal behavior among Neanderthal groups. Proceedings of the National Academy of Sciences USA 108, 250–253 (2011).

[7] Gavrilets, S. Human origins and the transition from promiscuity to pair-bonding. Proceedings of the National Academy of Sciences USA 109, 9923–9928 (2012).

